# Entrainment to sleep spindles reflects dissociable patterns of connectivity between cortex and basal ganglia

**DOI:** 10.1101/2022.02.06.479277

**Authors:** Aviv D. Mizrahi-Kliger, Alexander Kaplan, Zvi Israel, Hagai Bergman

## Abstract

Communication between the basal ganglia (BG) and cortex is crucial for behavior as it allows learning through external reinforcement. Non-REM sleep benefits learning in the corticostriatal system through the sleep spindle-associated reactivation of previously active neuronal ensembles and the subsequent modification of synaptic weights. However, how sleep spindles coordinate cross-region spiking, and whether spindle-driven reactivation occurs in other BG structures, remains unknown. We recorded field potentials (FP) and spiking activity in cortex and BG during sleep in two non-human primates immediately following a task that involved the learning of new cue-reward contingencies. FP sleep spindles were widespread in the BG, and they were similar to cortical spindles in morphology, spectral content and response to learning prior to sleep. Further, BG FP spindles were concordant with EEG spindles and associated with increased cortico-BG correlation. However, spindles across the BG differed markedly in their entrainment of local spiking. The spiking activity of striatal projection neurons exhibited consistent phase locking to striatal FP spindles and EEG spindles, producing phase windows of peaked cross-region spindling. In contrast, in the subthalamic nucleus (STN), which like the striatum receives substantial thalamocortical input, and in BG nuclei downstream to the striatum and STN, neuronal firing was not entrained to either local or EEG sleep spindles. These results dissociate striatal projection neurons from the rest of the BG, and suggest corticostriatal synapses as the main hub for offline communication between cortex and BG.

## Introduction

Sleep is believed to promote memory consolidation through offline reactivation of previously active neural ensembles (Oberto et al., 2022; Paller et al., 2021; Ramanathan et al., 2015). According to the “active system memory consolidation” theory, simultaneous reactivation of distributed brain networks during non-REM sleep facilitates plastic changes that consolidate initial labile representations into an enduring and retrievable format (Rasch and Born, 2013). Sleep spindles are ∼1 second long 11-15 Hz burst-like events, that are readily observed in cortical field potentials (FP) and EEG during non-REM sleep. Converging behavioral and mechanistic evidence suggest that offline reactivation of memory traces across distant brain networks is achieved through the interaction of slow oscillations and sleep spindles (Fernandez and Lüthi, 2020; Klinzing et al., 2019).

The basal ganglia (BG) are crucial for reinforcement learning, which is achieved through synaptic plasticity in corticostriatal synapses (Fisher et al., 2017). A role for sleep spindles in memory consolidation in the corticostriatal system is supported by several lines of evidence. Spindle amplitude was correlated with offline gains in performance in a motor learning task, as well as with corticostriatal BOLD activation signals (Barakat et al., 2013). An EEG-fMRI study employing a motor sequence learning task in humans found that the degree of striatal reactivations, time-locked to sleep spindles, was correlated with overnight task improvement (Fogel et al., 2017). Finally, a recent study in rodents associated sleep spindles with peak corticostriatal transmission and demonstrated their importance in modifying the spiking relationship between specific M1 and striatal neurons during skill learning (Lemke et al., 2021).

However, how sleep spindles coordinate cross-region communication between the cortex and BG remains unknown, as well as whether the striatum is the only BG nucleus that participates in offline modulation of connectivity strength with the cortex. To address these questions, we recorded cortical and BG spiking and field potentials (FP) along with multi-site EEG activity in two non-human primates (NHP). In addition to frontal cortex, we targeted the striatum and subthalamic nucleus (STN), which receive convergent thalamocortical innervation (Kincaid et al., 1998; Kolomiets et al., 2001), and two BG downstream structures, the GPe and GPi (Globus pallidus, external and internal parts) (Goldberg and Bergman, 2011).

Here, we report that although sleep spindles are prevalent throughout the BG, it is only in the striatum that spindling is associated with an increase in functional connectivity with cortex. While in the rest of the BG, spiking is disconnected from spindling, spiking activity of striatal projection neurons demonstrates consistent entrainment to both local and EEG spindling, and this entrainment reflects specific phase-windows of increased cross-region spindling.

## Results

Spectral analysis of basal ganglia (BG) field potentials (FP) during non-REM sleep in two non-human primates (NHP) revealed that spontaneous spindle-range activity was prominent in the striatum, STN, GPe and GPi as it was in the posterior dorsolateral frontal cortex (Fig. 1A). Applying conventional sleep spindle detection methods, we found that FP sleep spindles in the BG (see example in Fig. 1B) were similar to cortical spindles in duration (Fig. 1C), density (i.e., spindles per minute, Fig. 1D), amplitude (Fig. 1E) and spectral profile (Fig. 1F). We also found that like cortical FP sleep spindles, striatal spindles were modulated by local slow oscillations (SO), culminating in power (Fig. 1G) just before the FP SO peak. The same trend, albeit weaker, was observed in the STN, GPe and GPi.

**Figure 1.**
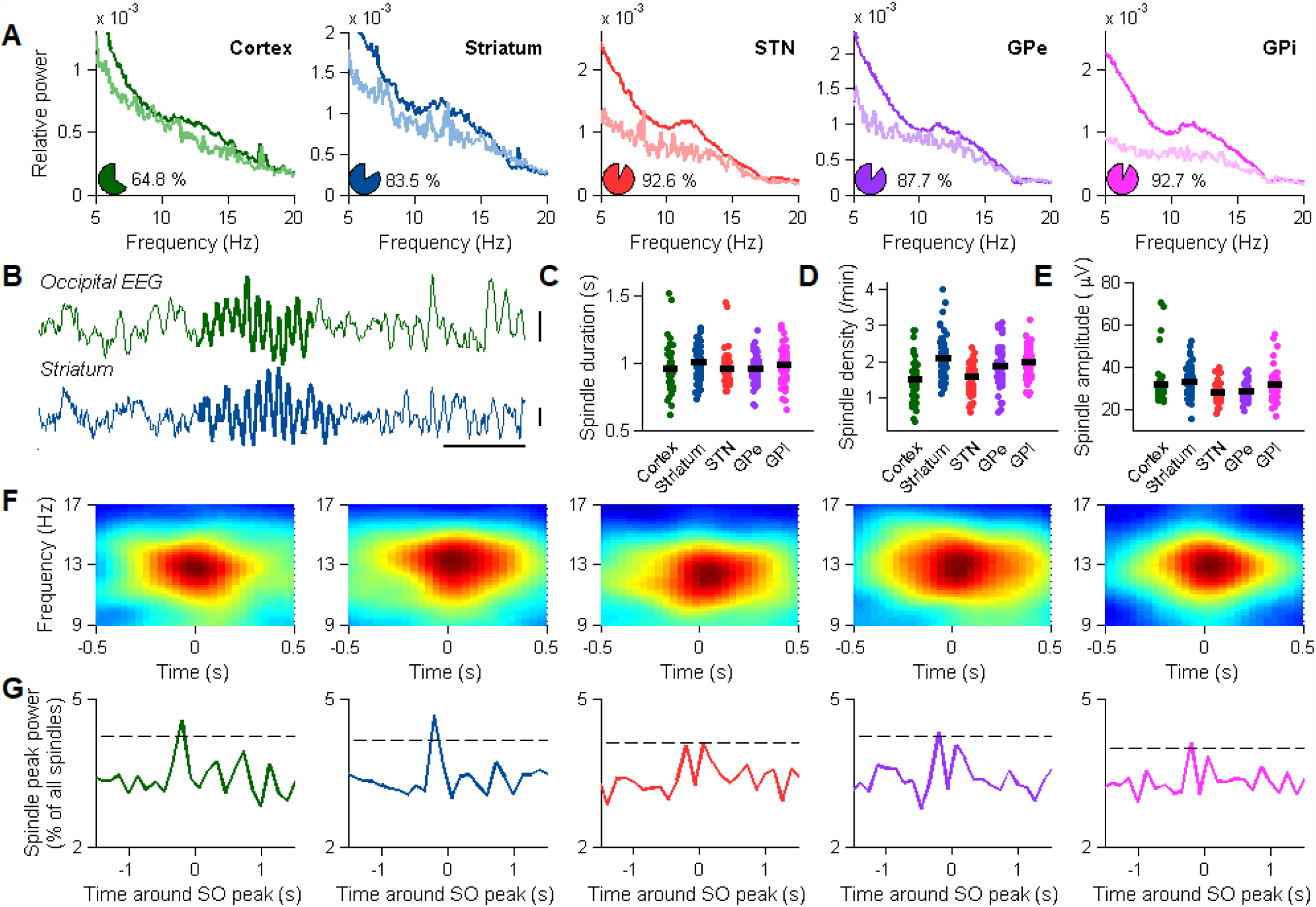
Sleep spindles are abundant in the BG during non-REM sleep, where they display similar characteristics as cortical spindles. (A) FP power spectra for non-REM sleep (dark) and wakefulness (light) in the frontal cortex (N=71 recording sites), striatum (N=103), STN (N=67), GPe (N=72) and GPi (N=82). Data is presented for sites that exhibited significant 10-17 Hz activity during non-REM sleep. Pie charts, the percentage of such sites, out of all recording sites. Unless stated otherwise, site numbers in the next subplots are as specified here. (B) Example ipsilateral occipital EEG (top) and striatal field potential (FP, bottom) showing concordant (synchronous) sleep spindles. Vertical bars, 50 μV, horizontal bar, 0.5 s. (C) Sleep spindle duration, cortex (N=2243 spindles), striatum (N=3390), STN (N=3007), GPe (N=4313) and GPi (N=4413). Horizontal black line represents the average. No significant difference, one-way ANOVA. (D) Sleep spindle density (number of spindles per minute). Horizontal black line represents the average. The cortex and STN were each significantly different from each of the striatum, GPe and GPi, one-way ANOVA and Tukey’s HSD procedure, p<0.05. (E) Sleep spindle mean amplitude. Horizontal black line represents the average. The STN and GPe were each significantly different from each of the cortex, striatum and GPi, one-way ANOVA and Tukey’s HSD procedure, p<0.05. (F) Average FP spectrograms for all sleep spindles recorded during non-REM sleep, cortex, striatum, STN, GPe and GPi. (G) FP sleep spindle power peaks relative to FP slow oscillation (SO) peaks, cortex, striatum, STN, GPe and GPi. Dashed line, p=0.01 confidence interval.

In addition to the cortical FP, we recorded the ipsilateral frontal and parieto-occipital EEG. The frequency characteristics of the EEG sleep spindles were similar to those recorded in the FP (Fig. S1A, S1E) as were sleep spindle duration (Fig. S1B), density (Fig. S1C) and amplitude (Fig. S1D). Intriguingly, unlike in human works (Andrillon et al., 2011; Purcell et al., 2017), we did not detect a difference in sleep spindle frequency between the frontal and parieto-occipital EEG (Fig. S1E). However, a recent work (Takeuchi et al., 2016) analyzing a large dataset of EEG sleep spindles in Japanese monkeys also reported only a small difference in spindle frequency between two anatomic locations similar to our frontal and parieto-occipital electrodes, as well as similar spindle durations, densities and amplitudes to those reported here. Also in accordance with results reported in that work, frontal and parieto-occipital EEG spindles were rarely concordant (Fig. S1F), and did not exhibit a consistent time lag (Fig. S1G).

The cortical FP and EEG spindles recorded here have mean frequencies of ∼13 Hz (Fig. 1F, Fig S1E), consistent with their characterization as fast sleep spindles (Mölle et al., 2011). Fast sleep spindles strongly relate to both declarative and non-declarative memory consolidation in cortical and subcortical brain regions (Muehlroth et al., 2019; Tamaki et al., 2008). The similar frequency characteristics of BG sleep spindles, along with their comparable duration and density, all indicate that cortical and BG spindles may reflect related phenomena. To examine whether this similarity could also be found at the functional level, we trained the animals to perform a task where they had to learn new cue-outcome contingencies, and examined the spindle response to learning. One to two hours prior to sleep onset, the animals performed a classical conditioning task in which they were presented with novel cues that were associated with either a positive (liquid food), negative (air puff) or neutral outcome, accompanied by a distinct auditory tone (Fig. 2A, and see methods). Along single blocks, during which each novel cue was presented ten times, the animals were able to learn the cue-outcome contingencies and perform anticipatory licking before the juice reward was given (Fig. 2B) (Kaplan et al., 2020).

**Figure 2.**
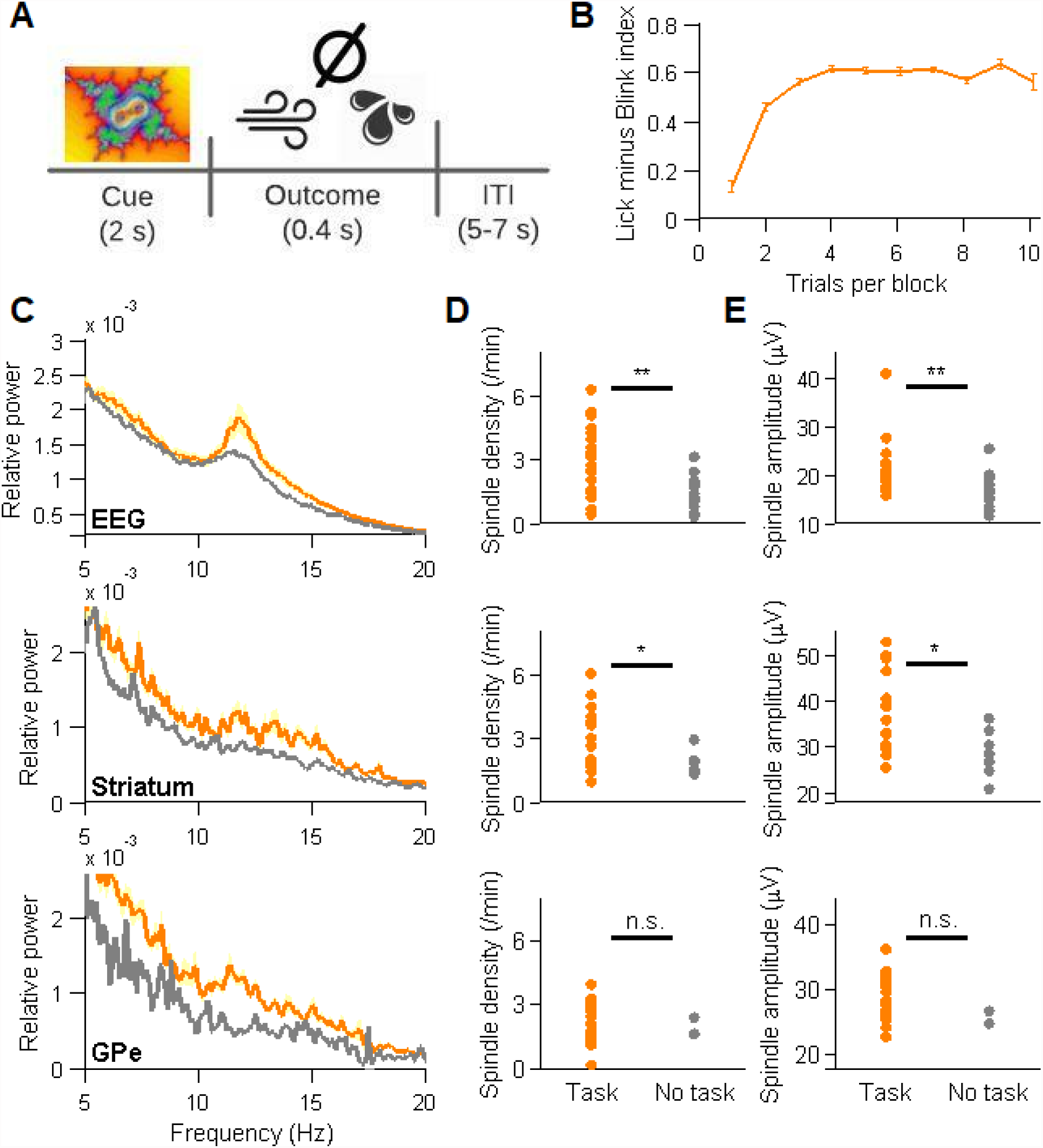
Learning of cue-reward contingencies prior to sleep is associated with an increase in sleep spindles in the EEG and striatal FP during early non-REM sleep. (A) A scheme of the behavioral task. Each trial, the animals were presented with a fractal cue followed by a juice reward, an air puff, or no reward. Novel cues were introduced each block, and presented along 10 trials. ITI, inter-trial interval. (B) Increase in Lick minus Blink index (reflecting anticipatory licking and decrease in blinking, which is associated with the aversive outcome. Range, 0-1, see methods) as a novel cue associated with a juice reward is learned along 10 trials (N=21 blocks over all recording days). Vertical bars represent s.e.m. (C) Top, average frontal and parieto-occipital EEG power spectra during the first 30 minutes of non-REM sleep after sleep onset for nights where the animals performed a learning task prior to sleep (orange, N=21) vs. nights where there was no task (gray, N=20). Pale shading represents s.e.m. 11-15 Hz power, task vs. no task, p=0.0017, Mann-Whitney U test. Middle, striatal FP power spectrum, task (N=21 recording sites) vs. no task (N=10 recording sites). 11-15 Hz power, task vs. no task, p=0.015, Mann-Whitney U test. Bottom, same for GPe FP power spectrum, task (N=23 recording sites) vs. no task (N=5 recording sites). 11-15 Hz power, task vs. no task, p=0.002, Mann-Whitney U test. (D) Top, EEG sleep spindle density for the first 30 minutes of non-REM sleep after sleep onset (for nights and electrodes in which significant spindle activity was observed), averaged across frontal and parieto-occipital EEG electrodes, task vs. no task. **, p<0.001, Mann-Whitney U test. Middle, same for striatal FP. *, p<0.05, Mann-Whitney U test. Bottom, same for GPe FP, n.s., non-significant, Mann-Whitney U test. (E) Top, EEG sleep spindle amplitude for the first 30 minutes of non-REM sleep after sleep onset (for nights and electrodes in which significant spindle activity was observed), averaged across frontal and parieto-occipital EEG electrodes, task vs. no task. **, p<0.001, Mann-Whitney U test. Middle, same for striatal FP. *, p<0.05, Mann-Whitney U test. Bottom, same for GPe FP, n.s., non-significant, Mann-Whitney U test.

We analyzed EEG and BG FP sleep spindle power, density and amplitude in the first 30 minutes of non-REM sleep, comparing nights where a task was performed and nights where there was no task prior to sleep. The average frontal and parieto-occipital EEG power spectrum (ipsilateral to our BG targets) showed an increase in sleep spindle power after the behavioral task (Fig. 2C, top row). Additionally, an increased sleep spindle density (Fig. 2D, top row) and amplitude (Fig. 2E, top row) were observed, in nights with a task vs. nights with no task, consistent with previous human results (Gais et al., 2002; Nishida and Walker, 2007). Sleep spindles were overall similar in their duration and spectral characteristics after a behavioral task and in nights where it was not performed (Fig. S2). Similar results were found when all EEG electrodes, and not only the ipsilateral ones, were analyzed. Strikingly, the striatal FP also exhibited an increase in spindle power (Fig. 2C, middle row), density (Fig. 2D, middle row) and amplitude (Fig. 2E, middle row) following the behavioral task. For GPe, spindle range power in the first 30 minutes of non-REM sleep was indeed increased (Fig. 2C, bottom row), but the behavioral task did not affect the spindle density or amplitude (Fig. 2D-E, bottom row. Early non-REM sleep data from nights with and without a task was available only for GPe, and not for GPi/STN). Thus, striatal and EEG sleep spindles are functionally similar in their response to learning.

The presence of cortical-like sleep spindles in the BG does not in itself indicate that the cortex and the BG spindles are related. However, when we recorded BG FP activity simultaneously with the ipsilateral EEG we found that non-REM sleep spindles in the BG tended to be concordant with EEG sleep spindles (meaning that if a spindle was detected in the striatum, there was a greater probability for a concurrent EEG spindle, compared to simulated data of two independent spindling processes, Fig. 1B and Fig. 3A). Further, within concordant spindles, the degree of overlap in duration with EEG sleep spindles was consistently higher than that expected by chance (Fig. 3B).

**Figure 3.**
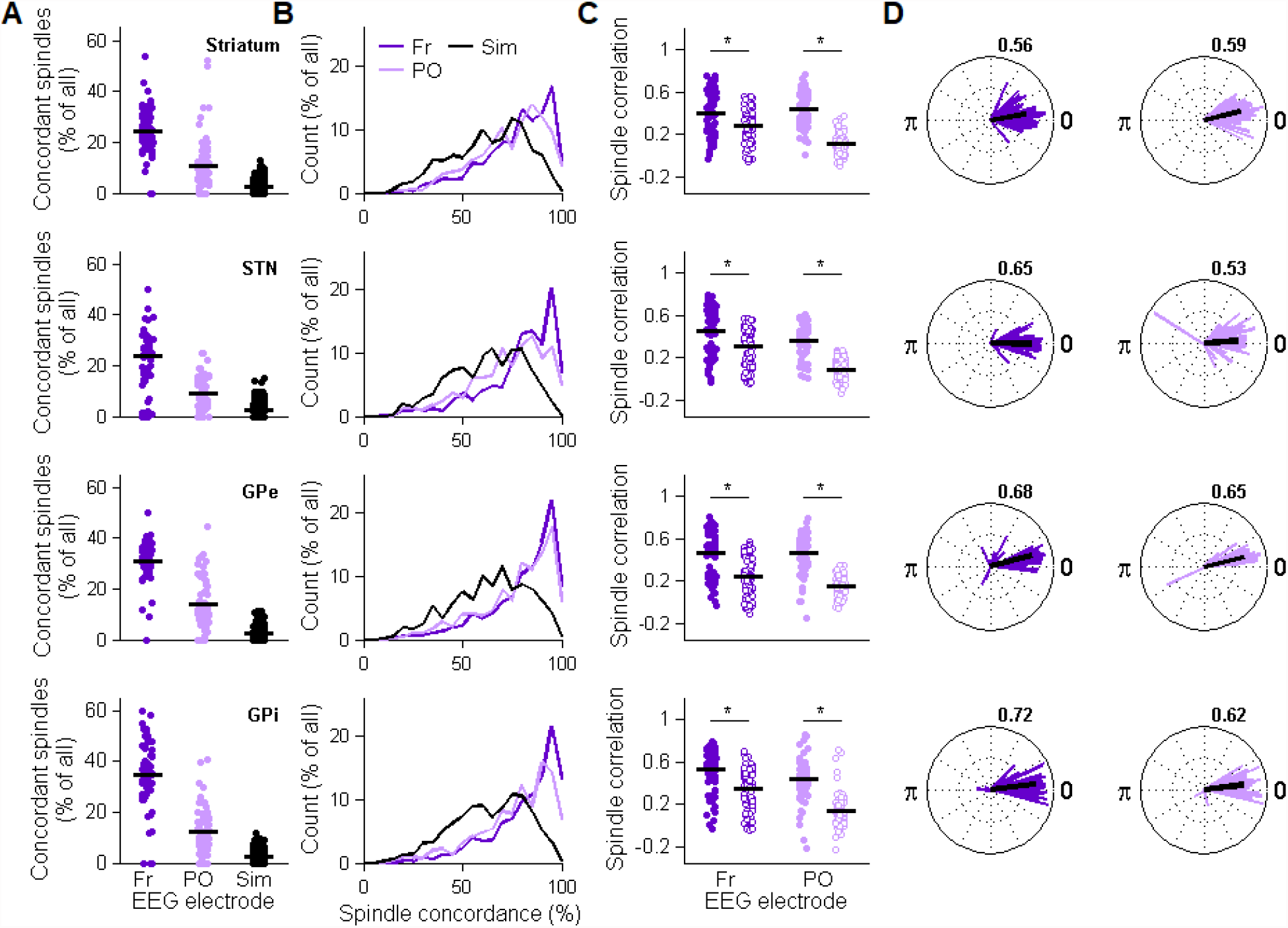
Sleep spindles are concordant between cortex and BG, and are associated with increased cortico-BG correlation. (A) Fraction of BG FP sleep spindles which are concordant (i.e., overlap) with EEG spindles, for ipsilateral EEG electrodes. Fr, frontal; PO, Parieto-occipital. Each point represents one recording site. Black, same for simulated FP and EEG sleep spindle data where the timing of spindles is random. Horizontal black line represents the average. Fr or PO vs. Sim, p<10^−4^ for all structures, Mann-Whitney U test. For all subplots, panels refer to (top to bottom) striatum, STN, GPe and GPi. (B) The degree of concordance for concordant EEG and FP spindles (in percent of the average EEG and FP spindle durations), for the two ipsilateral EEG electrodes, for all recording sites. Black, same for simulated data. Mean spindle concordance, Fr or PO vs. Sim, p<10^−4^ for all structures, Mann-Whitney U test. Color scheme as in (A). (C) FP-EEG correlation during sleep spindles (left) vs. during non-spindle segments of equal lengths (right), for the frontal and parieto-occipital EEG electrodes. Horizontal black line represents the average. *, p<5×10^−3^, Mann-Whitney U test. (D) Phase delays for simultaneously recorded sleep spindles in the BG FP and ipsilateral EEG. Each vector represents a single pair of FP and EEG recordings. The vector direction indicates the mean phase, calculated across all spindle pairs. The vector length indicates the phase lock index (PLI), i.e., the variability across all spindle pairs (0, maximal variability; 1, no variability). Horizontal black line represents the average. Population-average PLI is given in the top right.

BG sleep spindles were not only associated with concordant EEG spindling per se, but the overall cortico-BG functional connectivity was also increased. Sleep spindles were associated with increased correlation between the BG FP and the ipsilateral frontal and parieto-occipital EEG (relative to shuffled non-spindle data, Fig. 3C). To rule out the possibility that BG FP spindles represent volume conducted spindles from the cortex, we evaluated the distributions of delays between the waveforms of pairs of concordant EEG and FP spindles. The distributions appropriate for the volume conduction scenario would be strongly biased towards short positive delays. In our dataset, BG to ipsilateral frontal and parieto-occipital delays were both positive and negative, arguing against a strictly unidirectional propagation of sleep spindles in either way (Fig. 3D).

Cortical spindles entrain local spiking to specific spindle phases, and this has been suggested to support neural plasticity during non-REM sleep (Andrillon et al., 2011; Dickey et al., 2021). To examine whether local sleep spindles modulate single neuron spiking in the BG, we analyzed the spiking activity of BG projection neurons in relation to FP sleep spindles recorded in the same sites. We recorded the spikes and FP of striatal medium spiny neurons (MSNs), STN neurons, and GPe and GPi high-frequency discharging neurons. The occurrence of sleep spindles did not affect gross firing rates in either cortex or BG (Fig. S3). However, sleep spindles had a pronounced effect on spike timing in some, but not in all, nodes of the cortico-BG network. In the frontal cortex, neuronal spiking was modulated by spindle activity and locked to the troughs of the spindle waveform (i.e., a phase of π radians. Fig. 4A-D, top row). Similarly, sleep spindles in the striatal FP entrained MSN spiking to specific preferred phases which were concentrated around ∼1.67 π radians (Fig. 4A-D, second row). MSNs even showed significantly tighter locking to local spindles than cortical neurons, as indicated by higher phase locking indices (average PLI, 0.49 for MSNs vs. 0.25 for cortex, p=0.004, Mann-Whitney U test).

**Figure 4.**
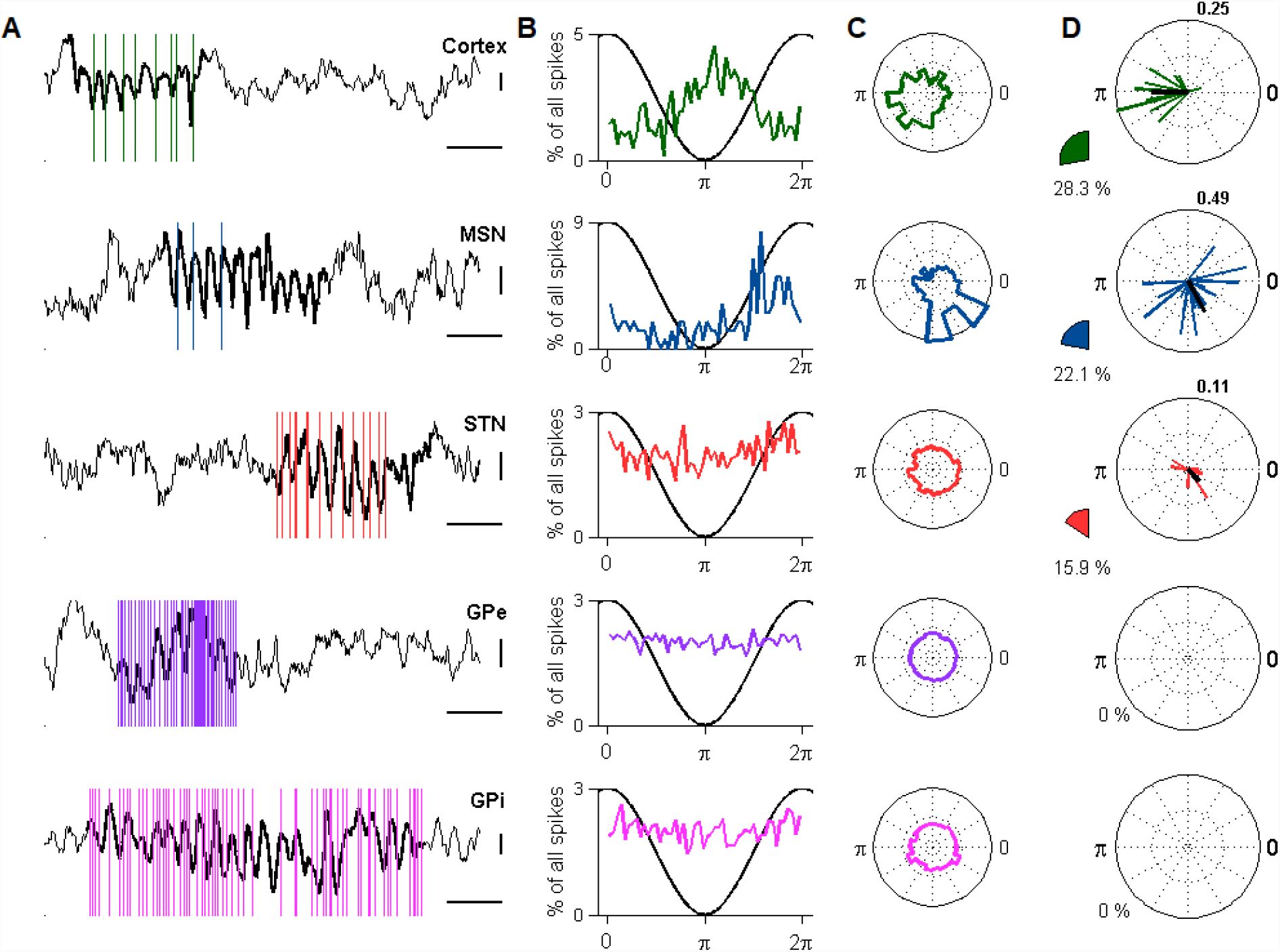
Sleep spindles entrain MSN spiking to specific phase windows but do not affect spiking in the rest of the BG. (A) Examples of neuronal spiking (vertical lines) relative to concurrent FP sleep spindles (bold) for the cortex, striatal MSNs, and STN, GPe and GPi neurons. Note cortical spike locking to spindle troughs and MSN locking to a later phase. Also note that STN, GPe and GPi spikes are not locked to a specific spindle phase. Vertical bars, 50 μV, horizontal bars, 0.25 s. (B) Distributions of the spindle phases of all spikes in the neurons depicted in (A). (C) Circular histograms depicting the preferred angles for the neurons in (A). (D) Entrainment of spontaneous spiking to FP sleep spindles in all cortical neurons (N=71), striatal MSNs (N=103), STN (N=67), GPe (N=72) and GPi neurons (N=82). Each neuron with significant and above-threshold (phase lock index, PLI, of 0.07) spike entrainment to sleep spindles is represented with a line, the angle of which indicates the preferred phase and the length of which represents the PLI. Black line, the average phase for all neurons. The population-average PLI is given in the top right. Notice that for GPe and GPi there was no spike locking to spindles. For visibility, cortex and STN vector lengths were multiplied by 2. Pie charts, proportion of neurons with significant and above-threshold spike entrainment to spindles, out of all neurons recorded in sites with significant FP spindle activity (p<0.05, Rayleigh’s test for non-uniformity).

Strikingly, whereas cortical and striatal spindling significantly entrained local spiking, spike discharge in the STN, GPe and GPi was apparently disconnected from spindling. In the STN, locking of spikes to FP spindles was relatively weak (Fig. 4A-D, third row). STN neurons were less likely to exhibit significant spike locking to sleep spindles (15.9% of neurons in comparison to 28.3% for cortex and 22.1% for MSNs). Further, the different STN neurons were not consistent in their preferred spindle phase, and the PLIs for STN neurons were significantly decreased relative to MSNs (average PLIs, 0.49 for MSNs, vs. 0.11 for STN, p=4.81×10^−6^, Mann-Whitney U test), suggesting a stronger MSN spike locking to spindles. For the GPe and GPi (Fig. 4A-D, bottom rows), no neuron produced significant above-threshold spike locking to sleep spindles.

The finding that cortical and striatal spindles are correlated, and separated, on average, by a short delay (Fig. 3D), suggests that striatal spike entrainment may not be locally bound. Indeed, MSN spiking was phase-locked not only to local spindling but also to concordant EEG spindles (Fig. 5A). In contrast, consistent with their lack of entrainment by local spindles, STN, GPe and GPi neurons showed weak to absent locking to EEG sleep spindles (Fig. 5B). Thus, cortical neurons and MSNs exhibit fairly consistent phase preferences to local spindles (Fig. 4D) and striatal spikes exhibit dual locking to cross-regional spindling in cortex and striatum. Generally, cortical activity is a major source of MSN excitatory input, and it is thus conceivable that spikes within striatal spindles derive from the firing of corticostriatal projection neurons. It is therefore possible that coordinated cross-region spindling may allow phase-specific spike transmission between cortex and striatum (Lemke et al., 2021).

**Figure 5.**
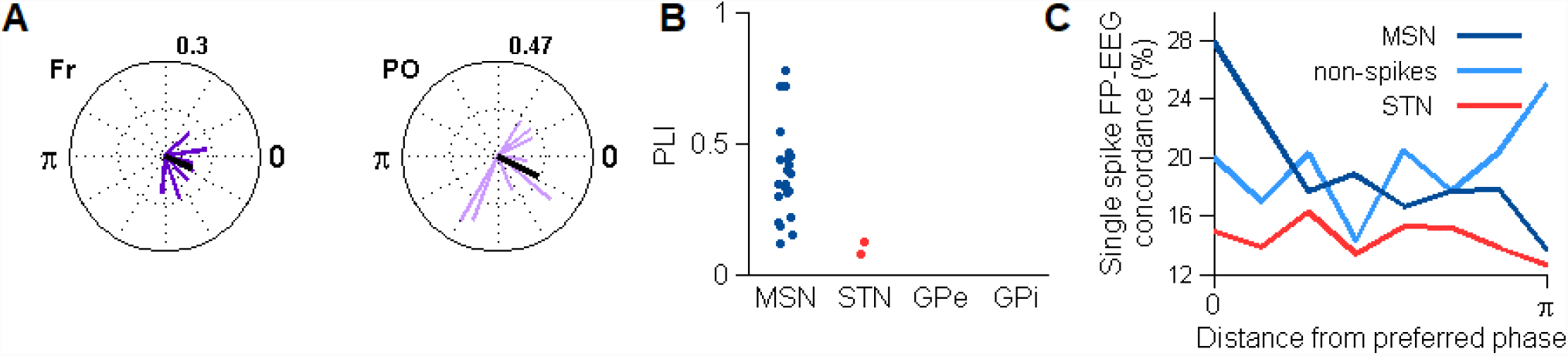
MSN spike locking to EEG and FP spindles reflects increased cross-region spindle concordance. (A) MSN spike phase locking to ipsilateral EEG sleep spindles (Fr, frontal. PO, parieto-occipital), with the corresponding mean angles and PLIs (top right, population-average PLI). (B) PLIs for all neurons exhibiting significant and above-threshold (0.07) spike locking to ipsilateral EEG spindles for MSNs, STN, GPe and GPi. Note lack of GPe or GPi locking to EEG spindles. (C) Single-spike FP-EEG concordance for MSNs, i.e. the probability that an MSN spike that was recorded within a striatal sleep spindle was also within an EEG sleep spindle, for spikes recorded in varying phase distances from the preferred locking phase (dark). Average over ipsilateral frontal and parieto-occipital EEG leads. Light trace represents samples within spindles that did not have spikes. Red trace represents STN spikes. Distance zero vs. any distance greater or equal than π/4, p<0.05, Chi-squared test. Same for non-spikes, n.s.

We evaluated this hypothesis by analyzing whether spikes that are close to the preferred phase are indeed associated with greater spindle concordance. Indeed, we found that in-spindle MSN spikes which are closer to the preferred phase were associated with higher concordance with EEG sleep spindles relative to spikes that were farther away from the preferred phase (Fig. 5C, dark trace). These changes might only be a function of spindle phase, rather than being associated with the actual spiking that took place in those phases. To correct for this possibility, we evaluated the FP-EEG concordance across the same range of spindle phases relative to the preferred phase, but for samples where no spikes were recorded. The spindle concordance rate did not vary across phases (Fig. 5C, light trace), indicating that an increase in cross-region spindling is associated with phase-specific spiking, rather than just spindle phase. Importantly, this effect was also specific to the striatum: spindle concordance was not affected by spiking phase in STN neurons which exhibited spike entrainment to spindles (Fig. 5C, red trace).

Further support for the hypothesis that spindle-entrained spiking in the striatum may represent cortical input comes from an indirect measurement of corticostriatal spike delays, locked to EEG sleep spindles. In our dataset, cortical and striatal spiking was not recorded simultaneously, but spiking in both structures was locked to ipsilateral frontal and parieto-occipital spindling. The mean phase delay between the preferred angles for cortical and striatal spiking was ∼26 degrees, which, for a 13-Hz oscillation, corresponds to a ∼6 ms delay, consistent with corticostriatal monosynaptic delay (Koralek et al., 2013).

## Discussion

Here, we show that sleep spindles prevail in the field potentials (FP) recorded in the BG during non-REM sleep and that they share statistical, morphological, spectral and functional properties with cortical/EEG sleep spindles. We also report that BG FP spindles are concordant with EEG spindles and associated with increased cortico-BG correlation. We proceed to demonstrate that striatal projection neurons (medium spiny neurons, MSNs) entrain local spiking to specific spindle phases, and that this locking reflects transient phase windows of increased cross-region spindling. This stands in sharp contrast to the rest of the BG, where spindles are prominent, but do not elicit any phase locked spiking.

Sleep spindles are generated through an intricate feedback excitation loop involving the thalamic reticular nucleus (TRN), and thalamocortical and corticothalamic projection neurons (Huguenard and McCormick, 2007; Nir et al., 2013). Spindle waveforms recorded in the scalp EEG are a result of synchronized 11-15 Hz burst spiking of underlying cortical neurons. However, deducing the origins of sleep spindles recorded in deeper brain structures is not straight-forward. Sleep spindles recorded in the BG FP could either be volume conducted (from cortex or thalamus) (Wennberg and Lozano, 2003) or reflect an actual cortico-BG or thalamo-BG spindle input (Haber and Calzavara, 2009; Steriade et al., 1984).

BG spindles were concordant with EEG sleep spindles (Fig. 3A-B) and were associated with increased cortico-BG connectivity (Fig. 3C), but to a moderate degree. Single spindle waveforms in BG could either lead or lag after EEG spindles (Fig. 3D). This is incompatible with a strong volume conductance effect, where concordance rates and correlations would be much higher, and single-waveform delay times would be highly homogenous and strictly positive. Alternatively, BG spindling could result from direct cortical inputs to either the striatum or the STN (Kincaid et al., 1998; Kolomiets et al., 2001). BG structures receive inputs from a range of cortical areas, and sleep spindles have different phases across the cortex (Nir et al., 2011; Takeuchi et al., 2016), so it is plausible that different BG spindles could be concordant with frontal or parieto-occipital spindles only for some of the time (Fig. 3A-B), span a range of correlations with them (Fig. 3C), and either lead or lag behind them (Fig. 3D).

Striatal spindle activity may also be the result of striatum-projecting neurons of the intralaminar thalamic nuclei (Galvan and Smith, 2011), which are among the major targets of the TRN (Steriade et al., 1984). The intralaminar nuclei are topographically organized such that the same regions project to striatal and cortical targets that are themselves functionally connected, i.e., involved in similar behavioral tasks (Haber and Calzavara, 2009). Given that functional rules govern also the anatomy of corticostriatal projections (Hooks et al., 2018) it is conceivable that cortical, thalamic and striatal areas that are active together during wakefulness will also spindle together during sleep. However, projections from the intralaminar nuclei also innervate the STN (Féger et al., 1994; Sadikot et al., 1992), so this possible route for sleep spindle propagation is not unique to the striatum..

Our core finding, that sleep spindles uniquely entrain striatal MSN spiking (Fig. 4), stresses the functional significance of BG spindles and differentiates MSNs (that make up more than 85% of striatal neurons, and are the only projection neurons of the striatum (Kita and Kitai, 1988)) from other BG neurons. The presence of consistent spike locking to specific spindle phases in the cortex and striatum (Fig. 4A-D, first and second rows) is compatible with a setting in which cortical action potentials are instigated in specific spindle phases and transmitted from cortex to striatum. Further support comes from our finding that the preferred phase for spindling in MSNs is associated with higher cross-region spindle concordance, relative to other spindle phases, in a way that is strictly spiking-dependent (Fig. 5C). The similar entrainment of MSN spiking to local and frontal/parieto-occipital spindles (Fig. 4D and 5A) also supports an MSN role in cross-region spindling. Indeed, sleep spindles were recently shown to facilitate corticostriatal transmission between M1 and dorsolateral striatum in rodents (Lemke et al., 2021).

Unlike MSNs and cortical neurons, STN, GPe and GPi spike locking to sleep spindles was weak to absent. What sets the striatum apart from the STN, GPe and GPi? The difference may be related to locking to slow oscillations (SO). In cortical recordings, sleep spindles are locked to up states of slow oscillations, and this locking has a wide functional importance (Fernandez and Lüthi, 2020). In our dataset, sleep spindles exhibited significant locking to FP SOs in the cortex and striatum but less so in the STN, GPe and GPi. SO locking may be important for spike alignment between distant brain regions, like cortex and striatum, and its absence in downstream BG structures may be detrimental for such alignment. Further, GPe and GPi neurons (and to a lesser degree, STN neurons) receive substantial GABAergic, mostly desynchronized, input from other GPe/GPi neurons (Goldberg and Bergman, 2011). This may disrupt the emergence of a population level spindle-aligned activity.

The dissociation between striatum and STN, GPe and GPi in spindle entrainment of spiking is supported by anatomical and functional considerations and resonated by findings from waking physiology. Striatal MSNs provide the majority of neural input to the downstream structures of the BG, but this input is inhibitory and relatively sparse (Bergman, 2021). The STN, similarly to the striatum, receives cortical and thalamic input, but provides substantial glutamatergic innervation to GPe and GPi (Kita and Kitai, 1987). This strong excitatory input is a powerful determinant of downstream BG activity during both wakefulness (Deffains et al., 2016; Schmidt and Berke, 2017) and sleep, as we recently demonstrated in a non-human primate model of Parkinson’s disease (Mizrahi-Kliger et al., 2020).

If indeed sleep spindles are positioned to allow precisely-timed recruitment of cortical cells and their respective striatal targets, such a mechanism could allow the spindle-instigated reactivation of cortical and striatal neuronal ensembles that were possibly active during a preceding waking period (Oberto et al., 2022). Such synchronous activation could strengthen their synaptic connectivity and underlie offline learning. Our observation that spindle range activity and sleep spindle density were elevated in the scalp EEG as well as in the striatal FP shortly after the completion of a task that required the learning of cue-outcome pairs (Fig. 2C-E) speaks in favor of a role for sleep spindle activity in the corticostriatal system in offline memory processing.

The current results supporting spindle-related offline increases in corticostriatal connectivity are in line with previous studies. EEG-fMRI works demonstrated that sleep spindles were associated with cortical and striatal reactivations during non-REM sleep and correlated with the degree of overnight improvement in a motor learning task (Boutin and Doyon, 2020; Boutin et al., 2018; Fogel et al., 2017). Electrophysiological work in rodents has reported the replay of task-related activity in motor cortex, linked to the coincidence of SOs and bursts of spindle activity, and implicated these replay events with offline improvements in a skilled upper limb task (Ramanathan et al., 2015). Finally, a recent work showed that striatal reactivations during sleep spindles reflected cortical input (Lemke et al., 2021).

These results establish that striatal medium spiny neurons and projection neurons from the rest of the BG represent two dissociable modes of connectivity with cortex during non-REM sleep. While sleep spindles are associated with increased functional connectivity between cortex and multiple BG nodes, it is only in the striatum that they entrain local spiking to specific spindle phases. These phase windows reflect transient increases in cross-region spindling, thus supporting corticostriatal connectivity and potentially spike transmission. If spike transmission within sleep spindles is taken to reflect reactivation of previous network activity that drives plasticity (Barakat et al., 2013; Fogel et al., 2017; Lemke et al., 2021), then our current results suggest that cortico-MSN synapses are the major cortico-BG synapses in which plasticity occurs, consistent with their being a prime target for dopaminergic reinforcement-based modulation (Bamford et al., 2018). All in all, our results suggest synchronized cross-region spindling as a main driver for synaptic plasticity in the corticostriatal system.

## Methods

All experimental protocols were conducted in accordance with the National Institute of Health Guide for Care and Use of Laboratory Animals and with the Hebrew University guidelines for the use and care of laboratory animals in research. The experiments were supervised by the institutional Animal Care and Use Committee of the Faculty of Medicine, the Hebrew University. The Hebrew University is an internationally accredited institute of the Association for Assessment and Accreditation of Laboratory Animal Care (AAALAC).

### Animals, sleep habituation and surgery

Data were obtained from two young adult female Vervet monkeys (*Chlorocebus aethiops sabaeus*, D and N) weighing ∼3.5 kg. The choice of non-human primates as a model for this current investigation was prompted by the similarities between non-human primates and humans in sleep architecture (Toth and Bhargava, 2013) and functional BG network organization (Hutchison and Everling, 2012).

The sleep recording routine was reported in a previous manuscript (Mizrahi-Kliger et al., 2018). Briefly, the monkeys were habituated to sleeping in a primate chair positioned in a dark, double-walled sound-attenuating room (IAC acoustics, IL, USA). The primate chair restrained the monkeys’ hand and body movements but otherwise allowed them to be in a position similar to their natural sleeping posture (Albert et al., 2011). The sleeping schedule was tailored to match the animals’ normal sleep hours, and recordings were made throughout the night (10-11 PM until 5-6 AM, 4-6 nights per week). Following sleep habituation, the monkeys underwent a surgical procedure for a 27×27 mm craniotomy and the implantation of a recording chamber, head holder, titanium screws for EEG recording, and trans-cranial ground screws (Mizrahi-Kliger et al., 2018). Recordings began after a postoperative recovery period of 5-7 days, during which an anatomical MRI scan was performed to estimate the chamber coordinates of the neuronal targets.

### Polysomnography and sleep staging

To determine the sleep stages throughout the night, the electroencephalogram (EEG), electrooculogram (EOG) and electromyogram (EMG) were recorded and the eye state (open/closed) was determined using continuous video recording. The scalp EEG was recorded from 5 locations: frontal (F3), central (Cz, C4) and posterior-occipital derivations (PO3 and PO4) (Sharbrough et al., 1991). EOG was recorded using disposable paired pre-gelled surface electrodes (Rhythmlink International, SC, USA). EMG was recorded using disposable paired (bipolar recording) subdermal needle electrodes (Rhythmlink International), inserted into the right trapezius muscle.

EEG, EOG and EMG signals were amplified with a gain of 20, filtered using a 0.075 Hz (2 pole) to 10 kHz (3 pole) Butterworth filter and sampled at 2,750 Hz by a 16-bit analog/digital converter. EEG and EOG were then digitally bandpass filtered using a zero-phase Butterworth filter in the range 0.1-35 Hz. EMG was bandpass filtered using a zero-phase Butterworth filter in the range of 10-100 Hz. For further analyses of the EEG spectra, the spectrum was divided by its total power post filtration to represent the relative power at 0.1-35 Hz. The positioning of electrodes, sampling and filtration followed the recommendations of the American Association for Sleep Medicine (AASM) Manual for Scoring of Sleep and Associated Events (Iber et al., 2007). The monkeys were digitally video-recorded at 50 frames/s using an infra-red camera (640×480 pixels, Norpix Inc., Canada) to assist sleep staging by detection of the eye open/closed state.

Sleep staging was performed using a semi-automatic staging algorithm (custom software) that used clustering of 10-second non-overlapping epochs, based on three features: the high/low frequencies EEG power ratio across all contacts, the root mean square (RMS) of the EMG signal and the eye-open fraction. Each 10-second epoch was represented as a point in a 3-dimensional feature space, creating three clusters corresponding to wakefulness, NREM sleep and REM sleep. All polysomnography and staging procedures were as detailed previously (Mizrahi-Kliger et al., 2018).

### Electrophysiological recordings and data collection

For the extracellular recordings, the monkeys’ heads were immobilized with a head holder and eight glass-coated tungsten microelectrodes were advanced separately (Electrode Positioning System, Alpha Omega Engineering, Israel) by two experimenters toward the target structures. Electrical signals were amplified with a gain of 20, filtered using a 0.075 Hz (2 pole) to 10 kHz (3 pole) Butterworth filter and sampled at 44 kHz by a 16-bit analog/digital converter. Spiking activity was sorted online using a template matching algorithm (SnR, Alpha Omega Engineering) after 300-6000 Hz filtration. BG neuronal assemblies were identified based on their stereotaxic coordinates according to MRI imaging and primate atlas data (Martin and Bowden, 2000) and real-time assessment of their electrophysiological features. Spiking and field potential (FP) were conducted only for unquestionably identified recording sites with stable recording quality (i.e., where single-neuron spiking yielded an average isolation score ≥0.85 (≥0.7 for STN neurons. The adjustment of the inclusion criteria for STN neurons was prompted by their relatively dense cellular arrangement, which made isolation difficult).

### Behavioral task

The behavioral task was discussed in detail previously (Kaplan et al., 2020). In short, we used a block design, where each block consisted of 30 trials. Each trial, a visual fractal cue was presented on a screen in front of the animal for two seconds, followed by either a positive outcome (juice reward), a negative outcome (an air puff) or a neutral outcome (no outcome). The reward, aversive and neutral trials were each associated with a single cue, so each block the animals were presented 3 different cues, 10 times each, in a pseudorandom order. Some of the blocks were overtrained, i.e., the animals knew the cues and the associated outcomes, and some were novel, i.e., for each block, a new series of 3 cues was presented and learned. Each novel or overtrained block could either be deterministic or probabilistic. In deterministic blocks, positive or negative outcomes were given at p=1 probability. In probabilistic blocks, positive or negative outcomes were given at p=0.5 probability. Blocks were grouped into sessions, each session comprising of 4 blocks: a deterministic overtrained block, a probabilistic overtrained block, a deterministic novel block and a probabilistic novel block, in a pseudorandom order. The animals typically performed 2-4 sessions before being allowed to sleep. Data used for analyses in this work was only taken from the deterministic novel block.

The animals’ task performance was assessed by monitoring their anticipatory licking and blinking, upon cue presentation and before the outcome was given. A digital video camera was used to record all sessions, and licking and blinking were automatically determined on the basis of the number of light and dark pixels in the mouth and eye area, using custom software (Mitelman et al., 2009).

For each reward trial, the number of licks/blinks was calculated by counting the number of licks/blinks from the time of cue onset until 1 s after cue onset. The lick minus blink index, indicative of the degree of learning for the rewarded cue, was constructed by subtracting the mean number of blinks from the mean number of licks and normalized to get real numbers between zero (minimal anticipatory licking) and one (maximal anticipatory licking).

### Power spectrum analysis

Each neuronal unit’s recording time was divided into 10-second epochs corresponding to the sleep staging epochs. We conducted a spectral analysis of the FP recorded in the vicinity of each of the analyzed neurons. The raw signal (hardware filtered between 0.075 Hz and 10 KHz, and sampled at 44 KHz) was digitally bandpass filtered using a zero-phase Butterworth filter (0.1-300 Hz, bandstop at 0.05 and 350 Hz for FP). For some of the recording sites, a high-power, very narrow band peak appeared at 16-17 Hz in the FP spectrum. Due to the extremely narrow frequency band characterizing it and its resemblance to higher frequency artefactual peaks (i.e. at 50 Hz), it was removed by a narrow notch filter.

Power spectra for FP signals were obtained using the *periodogram* Matlab function with a frequency resolution of 0.1 Hz. Different bin lengths, window functions and numbers of FFT points yielded similar results. The relative power at each frequency was then calculated by division of the entire spectrum by the overall power in the range of 0.1-50 Hz.

### Slow oscillation detection

Slow oscillation (SO) waveforms were detected in the 0.5-4 Hz filtered FP raw signal, as described in previous studies (Sela et al., 2016). Detected SO waveforms (i.e. between two consecutive negative-to-positive zero crossings) lasting between 0.25 second and 2 seconds were kept for further analysis. SO peaks and troughs were sorted according to their amplitude. Only peaks above the 70^th^ amplitude percentile were defined as SO peaks, and only troughs more negative than the 70^th^ amplitude percentile were defined as SO troughs. To verify that our detected events represent true slow oscillatory waveforms we examined whether SO peaks/troughs were associated with cortical spiking. As expected, FP SO troughs were associated with a significant increase in spiking, and the opposite was true for SO peaks.

### Sleep spindle detection and spike entrainment

Sleep spindle detection was performed on non-REM sleep EEG and FP data and was based on widely used conventional procedures (Sela et al., 2016). FP/EEG data was bandpass filtered (using a zero-phase Butterworth filter) between 10 and 17 Hz. This range was chosen because upon visual inspection of the power spectra, the spindle range occasionally surpassed 15 Hz. The Hilbert transform was then used to extract the instantaneous amplitude of the 10-17 Hz activity. Spindle detection was based on detecting events in time exhibiting high 10-17 Hz range activity. An initial detection threshold of 3 standard deviations above the mean was used to identify potential spindle events. A threshold of 0.5 standard deviation above the mean was then used to define sleep spindle start and end points. Due to its reliance on intrinsic variability, this detection method is relatively vulnerable to detecting sleep spindles where spindle activity is weak or non-existent. Thus, the detection algorithm was used only in those non-REM segments which exhibited significant 10-17 Hz activity in the power spectrum. A putative spindle event was identified as a sleep spindle only if it lasted 0.5 to 3 seconds, and only if it didn’t exhibit relatively high (4.5 standard deviations above the mean) instantaneous amplitude in a control 20-30 Hz range (this was done to exclude events with non-specific broadband power increases). Different frequency ranges and thresholds for amplitude and duration yielded similar results.

Spike entrainment to sleep spindles was based on detecting the exact sleep spindle phase corresponding to each spike (provided that it was detected within a sleep spindle), using the Hilbert transform of the 10-17 Hz filtered FP trace. For a given neuron, significant spike entrainment was based on Rayleigh’s test for non-uniformity for the distribution of all spike-related spindle phases.

### FP-EEG spindle connectivity measures

Spindle concordance was calculated for simultaneous FP and EEG recordings. For all detected sleep spindles in the FP, a concurrent EEG sleep spindle was defined as an EEG spindle starting 400 ms before or after the start of the FP spindle (Nir et al., 2011). Provided two concordant spindles were detected, the degree of concordance was defined as the duration of overlap time, as a percent of the average duration of both spindles. For sleep spindle FP-EEG concordance simulation, duration and inter-spindle intervals were randomly chosen from the actual distribution of per-structure FP spindle durations and inter-spindle intervals. The same was done for the EEG spindles basing on the frontal EEG spindle durations and inter-spindle intervals. Then, the above concordance measures were calculated in the same way as for recorded data.

For each FP sleep spindle, spindle correlation was defined as the correlation between the 4-40 Hz filtered FP and the concurrent EEG signals for the duration of the spindle. Notably, we did not require a sleep spindle to be present in the EEG to calculate correlation. The 0.1-4 Hz range was filtered out to avoid artifactual high or low correlations that derive from the presence of slow oscillations. Correlation were calculated identically for non-spindles segments, i.e. segments the same length of the actual sleep spindles that were randomly chosen from the same 10-second sleep epoch, provided they didn’t overlap a different detected sleep spindle.

Concordance was similarly calculated for single spikes. For each spike within each FP detected spindle, spike concordance was a binary variable reporting the presence (=1) or absence (=0) of a concurrent EEG spindle. Spike spindle concordance was calculated for all spikes within a certain range around the preferred entrainment spiking phase by calculating the average of the per-spike concordance values.

### Statistics

Analyses were conducted identically on the different neuronal assemblies. The data from the two monkeys were pooled since no significant differences were detected between them. The nonparametric Mann–Whitney U test was generally employed. A chi-squared test was used to compare proportions. Spindle duration, density and amplitude data between groups was analyzed using one-way ANOVA and Tukey’s Honestly Significant Difference (HSD) procedure as a post-hoc test, after the data was shown to distribute normally (using the Kolmogorov-Smirnov test). A threshold of 0.05, unless stated otherwise, was used to establish statistical significance. All statistical tests were two-tailed. Analyses and statistical computations were performed using Matlab 2013a (Mathworks, MA, USA). Data and code are available from the corresponding author.

## Acknowledgements

We thank Dr. Yaron Dagan and Dr. Tamar Ravins-Yaish for assistance with animal care. We thank Anatoly Shapochnikov, Dr. Hila Gabbay, Dr. Sharon Freeman and Dr. Uri Werner-Reiss for general assistance. We thank Dr. Marc Deffains for his assistance in neuronal recordings. Finally, we would like to thank Prof. Yuval Nir and his team members for their extensive assistance in planning the experiments, analyzing the data and finalizing the manuscript. This work was supported by grants from the Israel Science Foundation, Israel-China science foundation and the ReTune Germany Collaborative center TRR295 (to H.B.).

