## Supplementary figures and legends for "Entrainment to sleep spindles reflects dissociable patterns of connectivity between cortex and basal ganglia"

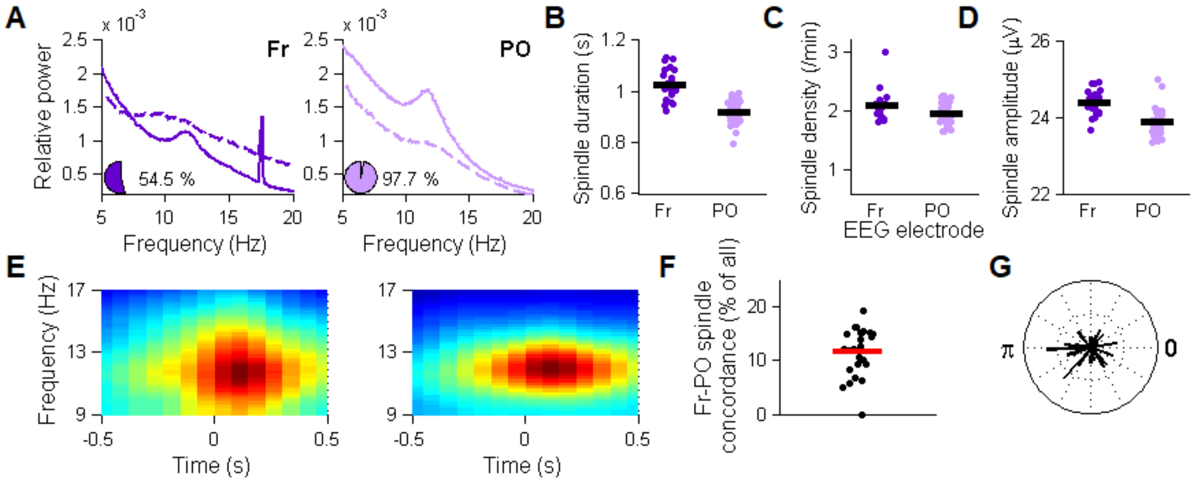


**Figure S1 – EEG spindle characteristics**

(A) Ipsilateral EEG power spectra for non-REM sleep (solid) and wakefulness (dashed) for all recording nights (N=44). Fr, frontal; PO, parieto-occipital. Pie charts, the percentage of nights exhibiting 10-17 Hz activity.

(B-D) Sleep spindle duration (B), density (C) and amplitude (D) for the nights and electrodes with significant 10-17 Hz activity. Horizontal black line represents the average.

(E) Average EEG spectrograms for all sleep spindles recorded during non-REM sleep, frontal (N=11418 spindles) and parieto-occipital (N=20795) electrodes. The EEG spectrograms look more granular relative to the FP spectrograms (Fig. 1) due to a lower sampling rate for EEG data.

(F) Percent of concordant spindles in the frontal and parieto-occipital electrodes, out of all frontal spindles. Horizontal red line represents the average conordance for all pairs.

(G) Phase delays for simultaneously recorded frontal and parieto-occipital sleep spindles. Each bar represents the mean delay of all concordant spindles in a single night. The radius of the unit circle corresponds to a PLI of 0.2. Across nights, no significant consistent delay was found.

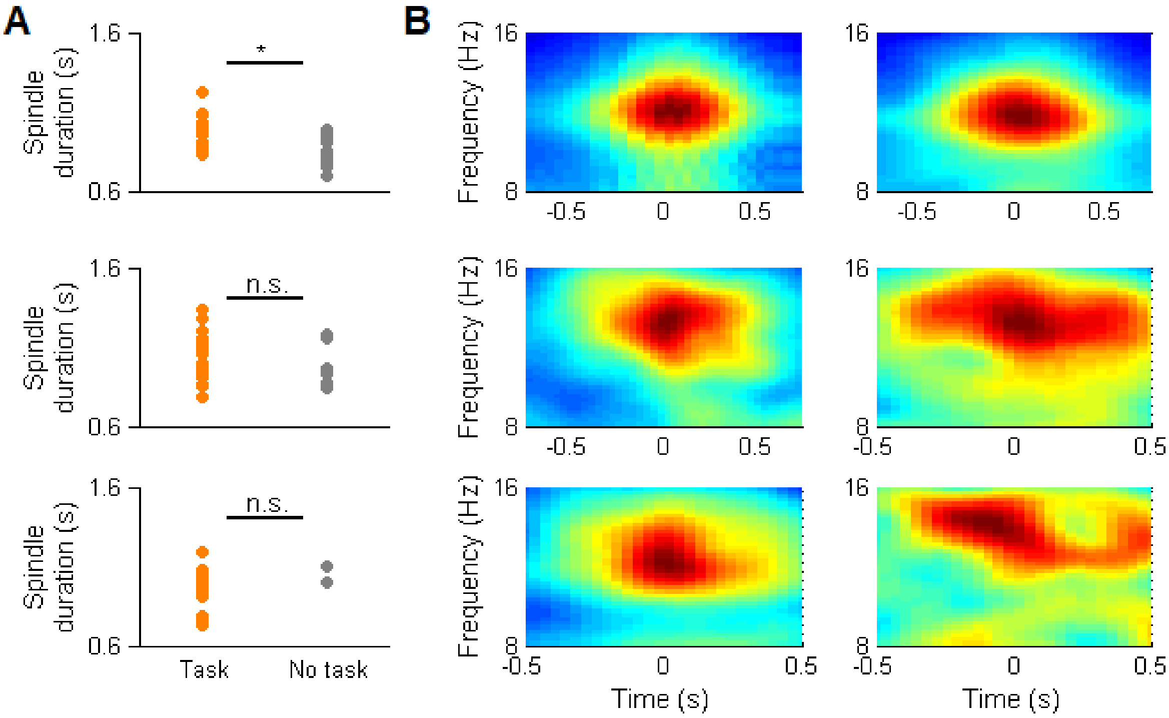


**Figure S2 – Spindle duration and spectral content are similar between nights with and without a behavioral task**

(A) Spindle duration during the first 30 minutes of non-REM sleep in nights with (left) and without (right) a behavioral task. Top, average ipsilateral frontal and parieto-occipital EEG; Middle, striatal FP; Bottom, GPe FP. *, p<0.005, Mann-Whitney U test. n.s., non-significant, Mann-Whitney U test.

(B) Spindle spectrograms for all spindles recorded during the first 30 minutes of non-REM sleep in nights with (right) and without (left) a behavioral task. Top, average ipsilateral frontal and parieto-occipital EEG; Middle, striatal FP; Bottom, GPe FP.

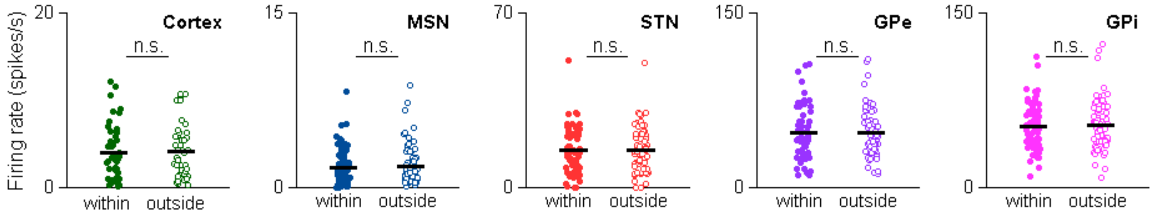


**Figure S3 – Firing rates are comparable within and outside of sleep spindles in the cortex and BG.**

Firing rates within (left) and outside of spindles (right) for cortical neurons, striatal MSNs and neurons of the STN, GPe and GPi. Horizontal black line represents the average. n.s., non significant, Mann-Whitney U test.
